# Proactive adjustments to cued gait perturbations in people with and without chronic stroke

**DOI:** 10.1101/2025.10.14.682416

**Authors:** Tara Cornwell, James M. Finley

## Abstract

Balance disturbances exist along a continuum from those that are fully unexpected to those that are predictable based on cues from the environment. When people experience predictable disturbances while walking, they may proactively adjust their gait to minimize losses of balance. However, one’s ability to effectively implement these proactive control strategies may be impaired after a stroke due to a combination of motor and cognitive impairments that result from brain lesions. Here, we used explicit audiovisual cues to characterize the proactive and reactive control strategies implemented by people with and without stroke during unexpected versus expected gait perturbations. Following unexpected treadmill accelerations, both groups had smaller margins of stability on the recovery step than during unperturbed walking. When we provided audiovisual cues specifying the impending perturbation step, people without stroke performed less leg and joint work, especially at the ankle, during the cued perturbations and increased their subsequent margins of stability by approximately 3 cm on the recovery step. However, people post-stroke did not make the same proactive adjustments. Instead, after any perturbation, they modified their stepping to maintain their center of mass position within their base of support, and this strategy remained unchanged with audiovisual cues. Our findings suggest that people post-stroke rely on a general control strategy rather than proactively modifying push-off work, even when given precise timing information about the impending gait perturbations.

## Introduction

People maintain balance while walking through a combination of proactive and reactive control strategies. Unexpected balance challenges, such as black ice, depend on reactive control adjustments informed by sensory feedback, whereas expected challenges, such as a visible change in terrain, allow for proactive control informed by awareness of an upcoming disturbance (1). Anticipatory postural adjustments, one example of proactive control, are characterized by changes in muscle activity based on knowledge of the timing, direction, or size of the expected movement or perturbation and have been examined during arm flexion tasks (2–4), gait initiation (5–7), and standing perturbations (8). Recent studies have used auditory cueing to test how knowledge of perturbation timing influences proactive control strategies during walking (9,10). When an auditory warning was provided three seconds before waist-pulls, treadmill-belt accelerations, or treadmill decelerations, young adults made no anticipatory changes to whole-body angular momentum or fore-aft margins of stability before or during the perturbation (10). It is possible that the young adults examined in this study did not generate proactive responses because they were confident in their ability to maintain balance using purely reactive strategies.

It stands to reason that populations with higher fall risk and impaired reactive control, such as older adults and people post-stroke (11–14), may increase their reliance on proactive balance control strategies. For example, compared to younger adults, older adults demonstrate more proactive adjustments when approaching an overground obstacle, progressively slowing and taking shorter steps (15), which increases the time available to take in visual information and prepare their control strategy. However, the ability to make proactive adjustments may be affected by stroke-related impairments, because compared to similarly-aged peers without stroke, people post-stroke produce less anticipatory muscle activity before experiencing an expected standing perturbation (16) and before initiating gait (7), which results in slower stepping (17), and before clearing a visible obstacle on a treadmill (18), which results in a higher failure rate (18,19). Proactive adjustments may be delayed or limited post-stroke due to somatosensory processing deficits (20), lower-extremity muscle weakness (21–23), abnormal muscle activation timing (24–26), or abnormal coordination (27,22,23). People post-stroke also frequently experience cognitive deficits in visual memory, executive function, and processing speed (28), which may affect their ability to understand audiovisual cues, plan an effective balance control strategy, and respond to cues promptly. Identifying how people post-stroke use timing cues to inform their control strategies could reveal how neurologic injury impairs proactive control during walking, possibly leading to falls.

The proactive balance adjustments made by older adults and people post-stroke while walking were recently examined by varying the predictability of perturbations using random- and regular-interval treadmill accelerations (29). Participants were not informed that perturbations would occur at fixed intervals in the regular condition, and, therefore, they needed to learn the regular pattern to proactively prepare for the perturbations. During regular perturbations, the group without stroke produced proactive adjustments that involved slowing the COM, but this strategy was not observed in their peers post-stroke (29). Given the need for participants to learn that the perturbations occurred regularly, it is possible that cognitive impairments limited the ability of people post-stroke to generate precise, proactive adjustments to their gait. In addition, the perturbations were applied unilaterally to the paretic side, but the control strategies may vary between the paretic and non-paretic sides due to the asymmetrical effects of a lesion.

Here, we examined how people with and without stroke used audiovisual cues to adjust proactive and reactive balance control strategies during single-belt treadmill accelerations. We varied the time between the cue and perturbation and the temporal precision of the cues to evaluate how control strategies changed as a function of the predictability of these perturbations. We quantified changes in fore-aft MOS and mechanical work performed by the legs to assess proactive adjustments and reactive responses to the perturbations. We hypothesized that people without stroke would use the audiovisual cues to proactively perform less positive work during the expected perturbation step relative to the no cue condition, resulting in a larger MOS during the recovery step. Because a stroke can cause a range of impairments in perception, sensorimotor processing, and control, we expected people post-stroke to have smaller reductions in positive work and smaller increases in MOS than the group without stroke, especially on their paretic side.

## Methods

### Participants

Based on an a priori power analysis using pilot data, we recruited fourteen chronic stroke survivors and 14 age-/sex-matched adults without stroke (Table 1) to complete this protocol. Several of the participants post-stroke reported using an ankle brace or orthotic (n=6), cane (n=2), or wheelchair (n=1) at least some of the time. We recruited participants from the University of Southern California Registry for Aging and Rehabilitation Evaluation (RARE) and the Healthy Minds Research Volunteer Registry. People post-stroke were included if they had a single stroke over six months prior and could walk unassisted for at least five minutes. Exclusion criteria for either group included musculoskeletal conditions that affect walking ability, uncontrolled hypertension, cognitive impairment (Montreal Cognitive Assessment score <19/30 (30)), or uncorrected visual impairment.

**Table 1.**
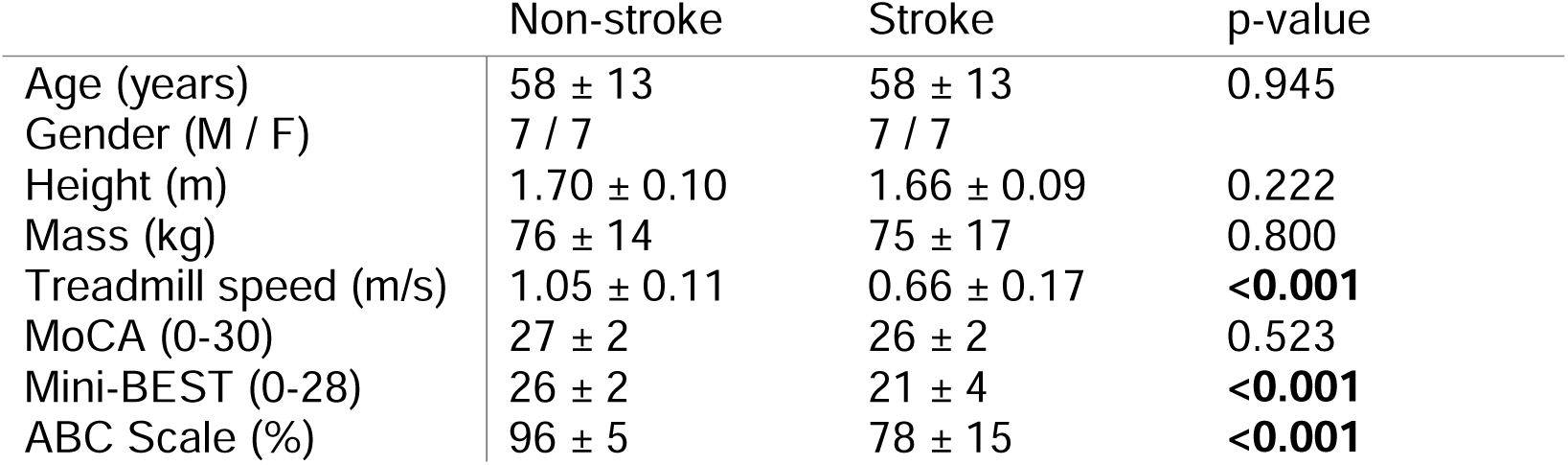
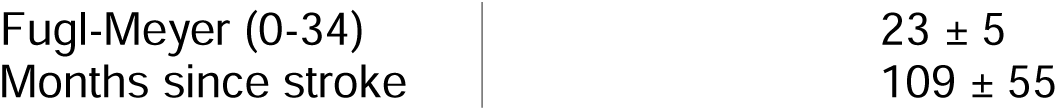
Group means ± standard deviations of demographic characteristics, clinical scores, and the p-values from Mann-Whitney U-tests evaluating group differences (bolded if p<0.05). MoCA: Montreal Cognitive Assessment, Mini-BEST: Mini Balance Evaluation Systems Test, ABC Scale: Activities-specific Balance Confidence Scale.

### Experimental protocol

We first conducted a set of clinical exams, including the Fugl-Meyer Assessment of lower-extremity impairment post-stroke, Montreal Cognitive Assessment (MoCA) to screen for cognitive impairment, Activities-specific Balance Confidence (ABC) scale to evaluate subjective balance confidence during daily tasks, Mini Balance Evaluations Systems Test (Mini-BEST) to evaluate dynamic balance, and 10-Meter Walk Test (10MWT).

After completing the clinical exams, participants completed six trials walking on an instrumented split-belt treadmill (Bertec, OH) at their self-selected walking speed (Figure 1A). They completed two baseline trials, during which they acclimated to the treadmill for one minute and then experienced perturbations (three accelerations per side). Finally, participants completed four trials with trip-like perturbations, during which a custom Python script accelerated one belt at 5 m/s^2^ up to 150% self-selected walking speed before returning to self-selected speed after the estimated stance time. We estimated swing and stance times as the average swing and stance times of the first 18 unperturbed steps per trial. Perturbations were triggered 250 ms before the estimated perturbation step to account for the time delays between sending the signal and accelerating the treadmill. We scaled perturbation magnitudes with walking speed to better normalize the relative effects of the perturbations across participants. Each perturbation trial consisted of 28 accelerations (14/side) every 12-22 steps with varied timing cues. The order of the four trials was randomized for every participant. The No-cue trial served as a reference condition. During the three trials with cues, participants received audiovisual cues (Figure 1B) and information about what the cues signified. All cues included an arrow denoting the side on which the upcoming perturbation would occur. A green light signified there was no upcoming perturbation. General cues (beep and yellow light) were triggered three to eight steps before the perturbation, Exact cues (beep and red light) were triggered two steps before, and Countdown cues (beeps with 3 in green, 2 in yellow, and 1 in red lights) were triggered six steps before. On average, the cues allowed participants post-stroke 3.1, 1.0, and 3.7 seconds and participants without stroke 2.6, 0.8, and 3.2 seconds to prepare before perturbations in the General-, Exact-, and Countdown-cue conditions. The cues varied in the time available to prepare before the perturbation and the precision of the information provided, allowing us to evaluate several aspects of proactive control. Participants wore a safety harness but did not have access to handrails or assistive devices throughout the trials.

**Figure 1.**
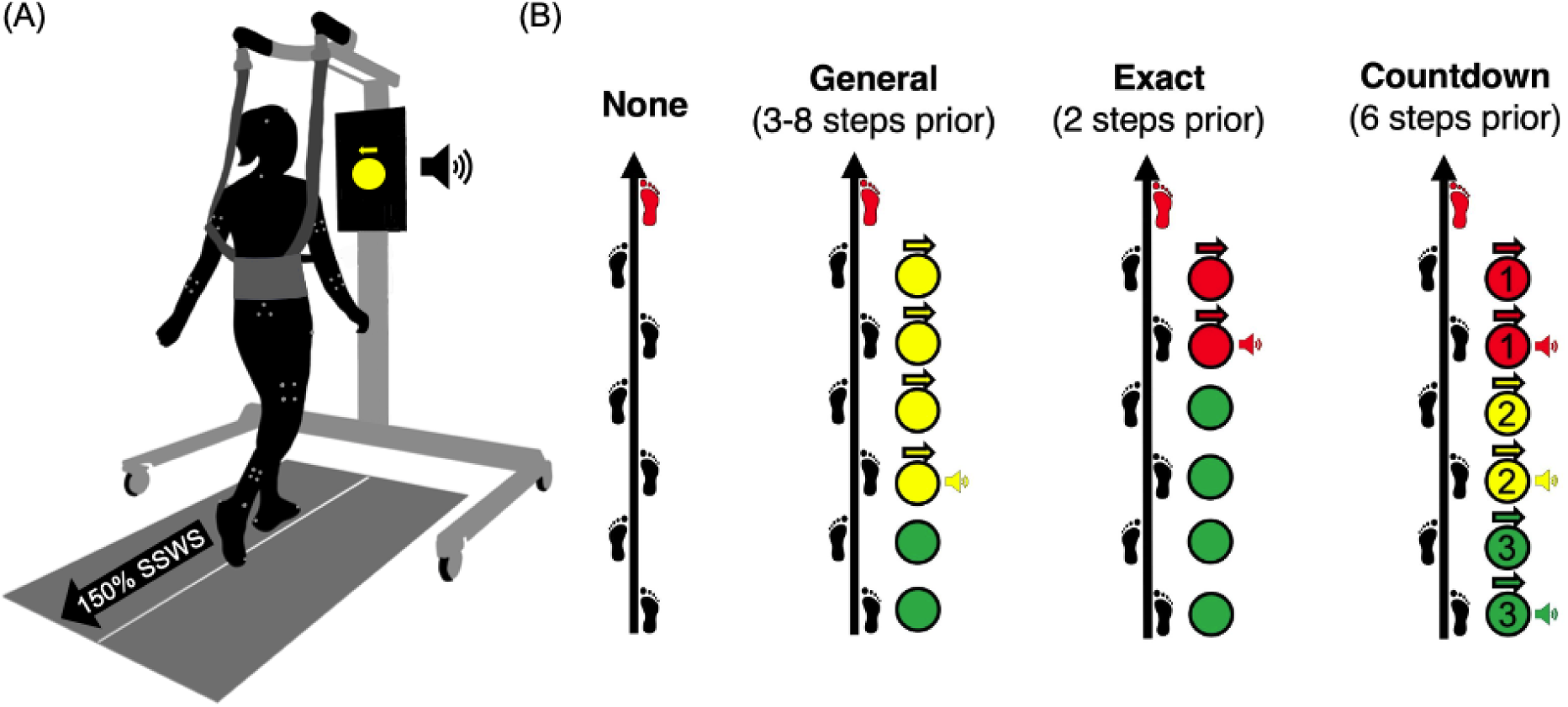
(A) The experimental setup with a participant wearing a full-body reflective marker set and safety harness on a split-belt treadmill while viewing cues on a monitor. (B) The four perturbation trials included No, General, Exact, and Countdown cues before the upcoming perturbation step (denoted by the red footstep). A green light indicated there were no upcoming perturbations. General cues (beep and yellow light) were triggered three to eight steps before the acceleration, Exact cues (beep and red light) were triggered two steps before the acceleration, and Countdown cues (beeps with 3 in green, 2 in yellow, and 1 in red lights) were triggered six, four, and two steps before the acceleration. All cues included an arrow to indicate the side of the upcoming perturbation.

### Data acquisition and processing

Participants wore 54 reflective markers on their feet, shanks, thighs, pelvis, torso, upper arms, forearms, and head. We used Qualisys Track Manager to track their positions using a 10-camera motion capture system at 100 Hz (Qualisys, Sweden). We also collected force data from the treadmill force plates at 1000 Hz.

We analyzed the data in Visual3D (HAS-Motion, Ontario, Canada) and MATLAB (MathWorks, Natick, MA). A custom Visual3D pipeline gap-filled (third-order polynomial with a maximum gap of 10 frames) and low-pass filtered (Butterworth with 6-Hz cutoff) the marker data. We then fit a 13-segment model to the processed marker data and used this model to estimate the position of the body’s center of mass (COM). We also used Visual3D to identify and visually confirm the following events: 1) heel strikes (HS) and toe offs as the maximum and minimum fore-aft positions of the heel and toe markers, respectively, 2) perturbation timing based on peak belt speeds, 3) the onset of visual cues recorded in an analog channel, and 4) “crossovers” when participants’ feet were placed on opposite belts. We removed force-based calculations during crossover events, which occurred, on average, four times per trial.

## Data analysis

We evaluated how people with and without a stroke modulated the fore-aft margin of stability (MOS) at HS, its components (COM speed and its distance from the edge of the base of support), positive mechanical leg work, and positive joint work (see Appendix) in response to treadmill perturbations and how these variables changed with audiovisual cues Because previous work has established that responses to perturbations may differ between the paretic and non-paretic sides (31), we separately evaluated our outcome measures depending on the perturbed limb in our group post-stroke. Therefore, for paretic perturbations, the perturbation step is taken by the paretic limb, and the subsequent recovery step is taken by the non-paretic limb, and the opposite is true for non-paretic perturbations.

### Fore-aft margin of stability

We evaluated the margin of stability (MOS), a common measure of dynamic stability, to test whether people made proactive modifications before the perturbation and effectively reduced the destabilizing effects following perturbations with versus without audiovisual cues. Because the perturbations were applied in the sagittal plane, we focused on the fore-aft MOS. The fore-aft MOS (Equation 1) was calculated as the anterior-posterior distance between the leading edge of the base of support (BOS) and the extrapolated COM position at HS, based on Hof’s inverted pendulum model (32).

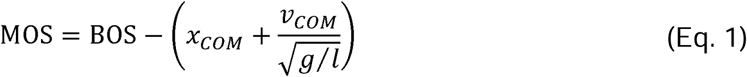

We defined the BOS as the fore-aft position of the toe marker, representing the anterior edge of the supporting area. The extrapolated COM was computed as the fore-aft COM position (x_COM_) adjusted by fore-aft COM speed relative to the treadmill belt (v_COM_), gravity (g), and leg length (l). Negative MOS values indicate that the extrapolated COM is anterior to the BOS, and a corrective step or adjustment is generally required to prevent a fall forward. We hypothesized that people, especially those without stroke, would use audiovisual cues to proactively increase their MOS at the HS of the perturbation step to reduce the destabilizing effects. Subsequently, if participants effectively updated their strategy with audiovisual cues and reduced the destabilizing effects of the perturbation, we hypothesized that people, especially those without stroke, would have a larger MOS at the recovery step HS.

Since the MOS is a function of foot placement relative to the COM and COM speed, changes in the MOS could result from changes in either or both variables. Therefore, we decomposed the MOS into its components and evaluate how each changed across cueing conditions. These components included the distance between the edge of the BOS and COM position (BOS-COM) and COM speed, both of which are evaluated at HS of the perturbation and recovery steps to capture changes to proactive control and subsequent stability, respectively. We calculated BOS-COM as the fore-aft distance between the front toe marker and COM and the fore-aft COM speed relative to the treadmill belt.

### Positive leg work

When walking, people perform mechanical work with their legs to advance their COM position from step to step. Prior work found that people proactively reduce positive mechanical leg work and reduce the destabilizing effects of expected treadmill perturbations (29). Therefore, when given cues, we expect people to proactively update their control strategy and perform less positive leg work during the pre-perturbation or perturbation steps to increase recovery step MOS, either by slowing the forward COM speed or reducing the forward displacement of the COM. We computed mechanical leg power (P_leg_) in Watts using the individual limbs method (33) as the sum of the power performed by the leg on the body and the treadmill (Equation 2).

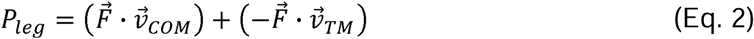

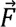 is the ground reaction force vector, 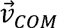 is the COM velocity vector, and 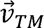 is the treadmill velocity vector relative to the lab coordinate system. We calculated positive work for each step by integrating the positive portions of the power curve and then normalized work by the subject’s mass, resulting in positive leg work in Joules per kilogram.

### Statistical analysis

Our primary aim was to understand how each outcome measure varied by group and cue condition. While adaptation to repeated perturbations was possible, we mitigated confounding effects of adaptation by evaluating the median of each measure calculated per trial. First, we determined if the effects of the perturbations on COM dynamics during the No-cue trial differed between groups by fitting a linear mixed-effects model for the change in median fore-aft COM speed at HS of the recovery step (Equation 3). The change in COM speed (Δv) was computed relative to unperturbed walking, captured during the first baseline trial. The model included an intercept (β_NS,No_), fixed effect of Leg (NS: non-stroke, P: paretic, NP: non-paretic), and random intercepts (b_i_) to account for variability between subjects and sides within the group post-stroke. If the results of an ANOVA yielded a significant fixed effect of Leg, we reported the coefficients and their p-values, which were Bonferroni corrected for the two comparisons.

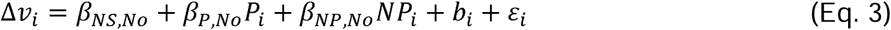

Here, i is the observation number (1:42) and ε is the error.

Then, we fit separate linear mixed-effects models (Equation 4) for the changes in the median fore-aft MOS, fore-aft COM speed, fore-aft BOS-COM distance, positive leg work, and positive joint work relative to baseline (Δy) with fixed effects of Leg (NS: non-stroke, P: paretic, NP: non-paretic), Cue (No, Gen: General, Ex: Exact, and Count: Countdown), and the interaction between Leg and Cue. Baseline values are the medians calculated during the one-minute, unperturbed walking trial. We considered changes in positive leg and joint work during the pre-perturbation step, all metrics during the perturbation step, and MOS, BOS-COM, and COM speed during the recovery step. The models included random intercepts (b_j_) to account for variability between subjects and sides within the group post-stroke. If the results of an ANOVA yielded significant fixed effects, we reported the coefficients and their p-values, which were Bonferroni corrected based on the number of comparisons (two for Leg, three for Cue, and six for Leg*Cue).

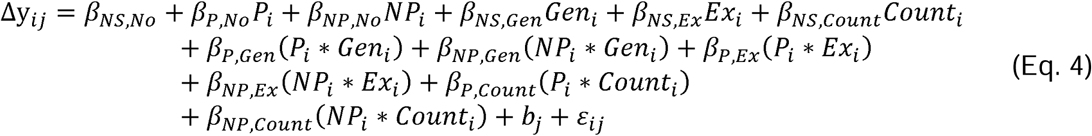

where i is the observation number (1:168), j is the leg (1:42), β is the coefficient per fixed effect, and ε is the error.

All statistical analyses were performed using MATLAB 2022b with an alpha level of 0.05. For every model, we plotted standardized residuals versus fitted values to check for heteroscedasticity, nonlinearity, and outliers. Outliers were identified if the standardized residual was outside three standard deviations, and the model was refit, excluding the outliers.

## Results

### Treadmill perturbations resulted in faster COM speeds in people with and without stroke

The treadmill perturbations caused an increase in fore-aft COM speed at the HS of the first recovery step for participants in both groups (Figure 2A). The perturbations in the No-cue condition increased COM speeds by 0.12 ± 0.01 m/s at the recovery step HS (Figure 2B) relative to unperturbed walking (Equation 3; β_NS,No_ p<0.001). There was no significant difference in the change in speed between groups (Leg p=0.254), indicating that the perturbation intensity was similar for participants with and without stroke.

**Figure 2.**
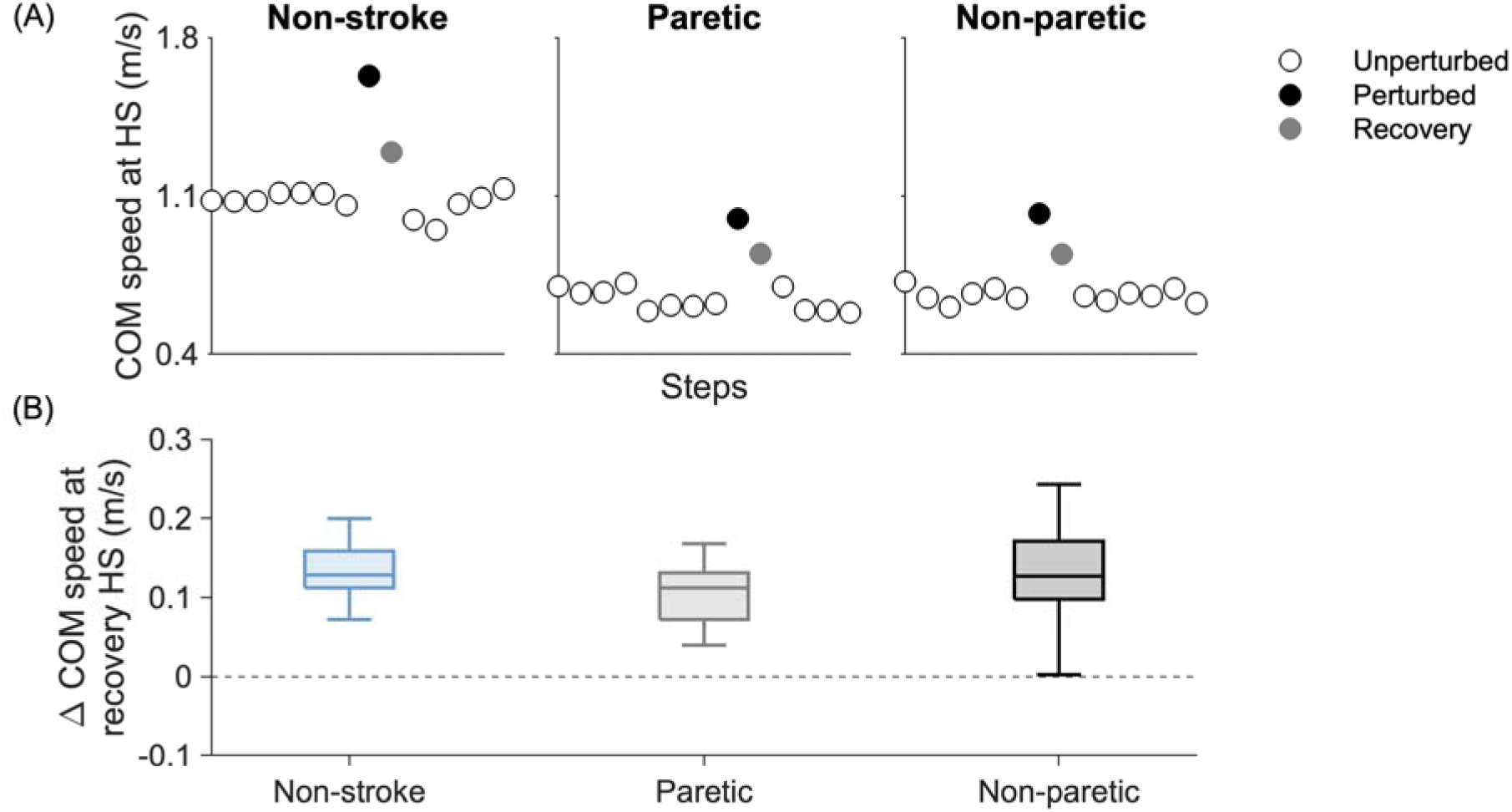
(A) The fore-aft center of mass (COM) speed at heel strike (HS) of unperturbed (white), perturbed (black), and the first recovery (gray) steps for representative participants without stroke (Non-stroke) and with chronic stroke when perturbed on their paretic (Paretic) or non-paretic (Non-paretic) side during the No-cue trial. (B) The change in COM speed at the recovery step HS after perturbations in the No-cue trial. The change was calculated relative to the average COM speed during unperturbed steps. Boxplot colors represent perturbations applied to the three legs: Non-stroke (blue), Paretic (gray), and Non-paretic (black). For paretic perturbations, the change in COM speed is computed as the change in speed during the non-paretic recovery step relative to non-paretic steps during the unperturbed baseline trial. Similarly, for non-paretic perturbations, the change in COM speed is computed as the change in speed during the paretic recovery step relative to paretic steps during the unperturbed baseline trial. The change in COM speed at the recovery step HS did not statistically differ between legs (p=0.254), suggesting similar perturbation intensity across groups. Three total outliers were omitted to maximize the visual size of the boxplots.

### Precise cues led to more positive MOS values during the recovery step, especially in people without stroke

Next, we tested whether people made proactive changes in the fore-aft MOS at the HS of the perturbation step (Figure 3A). The increase in belt speed during perturbations resulted in a general reduction in the MOS during the perturbation step relative to unperturbed walking. The MOS at the perturbation HS was 15.4 ± 0.7 cm smaller for the non-stroke group (β_NS,No_ p<0.001), 9.0 ± 1.1 cm smaller in the group post-stroke during paretic perturbations (β_P,No_ p<0.001), and 9.1 ± 1.1 cm smaller in the group post-stroke during non-paretic perturbations (β_NP,No_ p<0.001). However, there was no effect of cueing on the perturbation step MOS (Cue: p=0.095; Interaction between Leg and Cue: p=0.965). Thus, cues did not elicit proactive changes in MOS at the time of the perturbation.

**Figure 3.**
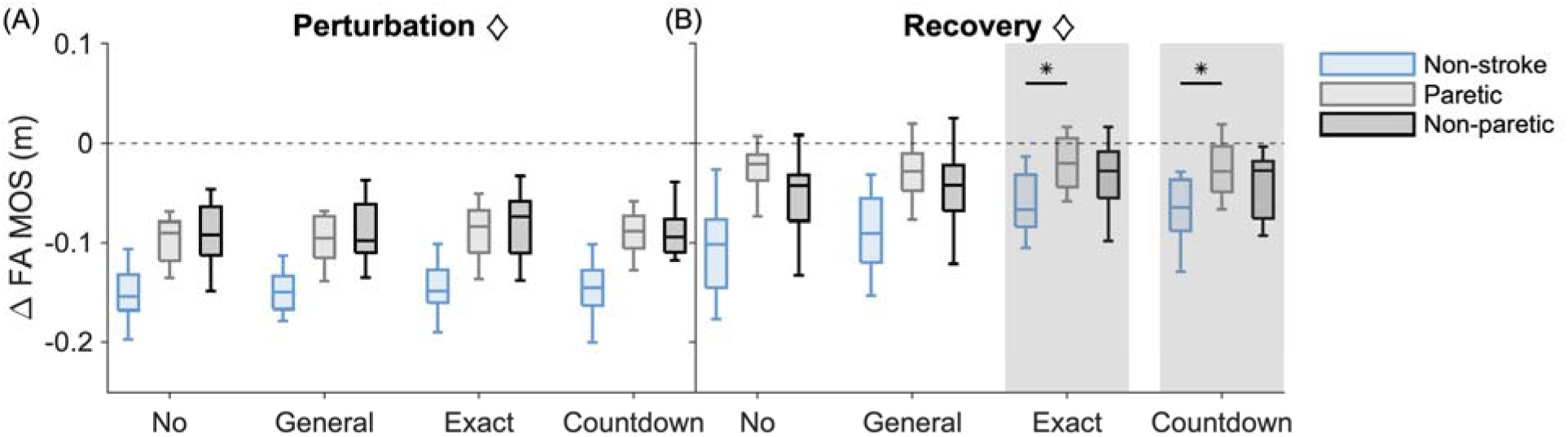
Change in the fore-aft (FA) margin of stability (MOS) during (A) perturbation and (B) recovery steps with No, General, Exact, and Countdown cues. The changes were calculated relative to unperturbed walking, with negative values indicating the extrapolated center of mass is more anterior relative to the leading edge of the base of support when compared with its position at unperturbed heel strikes. Boxplot colors represent perturbations to the Non-stroke (blue), Paretic (gray), and Non-paretic (black) limbs. Diamonds in the title indicate significant main effects of Leg. The results of post hoc analyses evaluating the main effect of Cue or its interaction with Leg are represented by vertical gray shading, indicating significant differences from the No-cue trial, and asterisks, indicating significant within-condition differences between the non-stroke and stroke groups. Five total outliers were omitted to maximize the visual size of the boxplots.

Next, we evaluated changes in the fore-aft MOS at HS of the recovery step to test if cues helped people improve stability (Figure 3B). For people without stroke, these MOS values remained 9.1 ± 0.9 cm smaller during the recovery step relative to unperturbed steps (β_NS,No_ p<0.001), but this difference between unperturbed and recovery step MOS was smaller in the group post-stroke after both paretic (β_P,No_ p<0.001) and non-paretic (β_NP,No_ p=0.002) perturbations. There was a significant effect of cueing on the difference between the recovery step MOS and unperturbed MOS, but it differed between groups (Cue: p<0.001; Interaction between Leg and Cue: p=0.015). Compared to the No-cue trial, people without stroke increased MOS at the recovery step HS by 2.9 ± 0.5 cm with Exact cues (β_NS,Ex_ p<0.001) and by 2.5 ± 0.5 cm with Countdown cues (β_NS,Count_ p<0.001) but did not change MOS with General cues (β_NS,Gen_ p=0.959). However, following paretic-leg perturbations, people post-stroke made smaller changes in recovery step MOS than their peers without stroke by 2.1 ± 0.7 cm with Exact cues (β_P,Exact_ p=0.032) and by 2.3 ± 0.7 cm with Countdown cues (β_P,Count_ p=0.012). Overall, more precise timing cues allowed people, especially those without stroke, to increase their recovery step MOS values.

We also evaluated changes in BOS-COM distance at the HS of perturbation (Figure 4A) and recovery (Figure 4B) steps separately from the calculation of MOS to understand the role of COM position in cue-related changes in stability. The BOS-COM distance on the perturbation step did not differ from unperturbed walking (β_NS,No_ p=0.785) or between groups (Leg: p=0.875). However, there was a significant effect of cueing on BOS-COM distance (Cue: p=0.024; Interaction between Leg and Cue: p=0.927) with people increasing BOS-COM distances on the perturbation step by 1.0 ± 0.4 cm with Exact cues (β_NS,Ex_ p=0.040) but not with General (β_NS,Gen_ p=1.000) or Countdown (β_NS,Count_ p=0.055) cues. These changes were small, suggesting that proactively adjusting their COM position within their BOS was not the main control strategy, which is consistent with the lack of changes in the MOS during the perturbation step.

**Figure 4.**
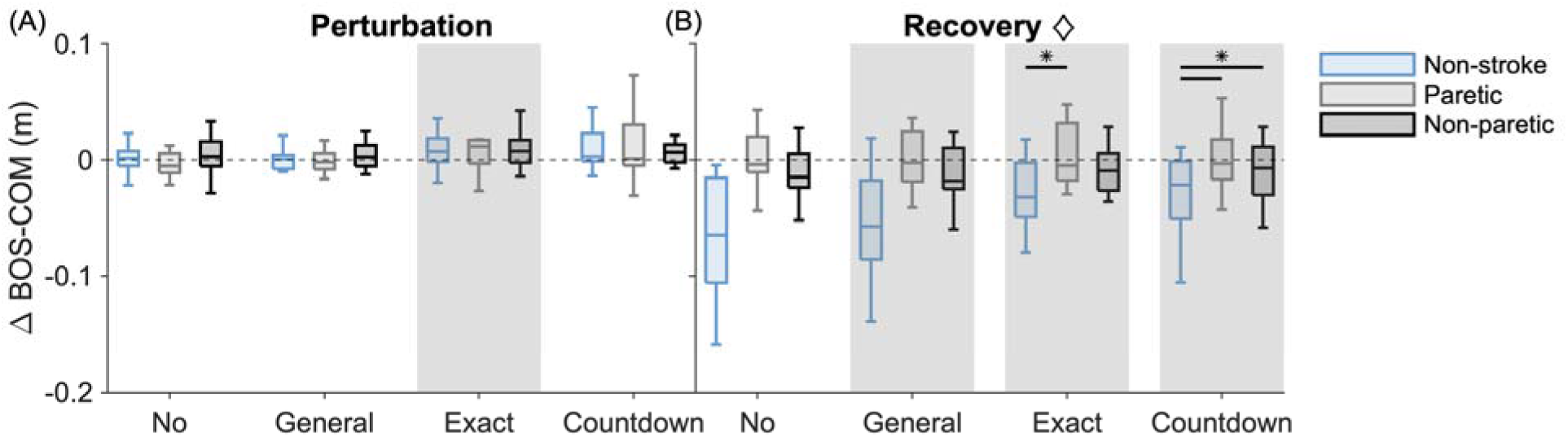
The change in the fore-aft distance between the edge of the base of support (BOS) and center of mass (COM) at heel strike of (A) perturbation and (B) recovery steps with No, General, Exact, and Countdown cues. The changes were calculated relative to unperturbed walking, with negative values indicating the COM is more anterior relative to the leading edge of the base of support when compared with its position at unperturbed heel strikes. Boxplot colors represent perturbations to the Non-stroke (blue), Paretic (gray), and Non-paretic (black) limbs. Diamonds in the title indicate significant main effects of Leg. The results of post hoc analyses evaluating the main effect of Cue or its interaction with Leg are represented by vertical gray shading, indicating significant differences from the No-cue trial, and asterisks, indicating significant within-trial differences between the non-stroke and stroke groups. Eleven total outliers were omitted to maximize the visual size of the boxplots.

After the perturbation in the No-cue trial, the BOS-COM distance on the first recovery step (Figure 4B) was 5.5 ± 0.8 cm smaller for people without stroke (β_NS,No_ p<0.001), 0.2 ± 1.1 cm larger for people post-stroke following paretic perturbations (β_P,No_ p<0.001), and 1.5 ± 1.1 cm smaller following for people post-stroke non-paretic perturbations (β_NP,No_ p<0.001) compared to unperturbed walking. Cueing significantly affected the size of the BOS-COM distance at the recovery step HS, but this effect differed between the groups (Cue: p<0.001; Interaction between Leg and Cue: p=0.017). For people without stroke, their BOS-COM distances on the recovery step were 1.3 ± 0.5 cm, 2.2 ± 0.5 cm, and 2.4 ± 0.5 cm less negative with General (β_NS,Gen_ p=0.025), Exact (β_NS,Ex_ p<0.001), and Countdown (β_NS,Count_ p<0.001) cues, respectively, than in the No-cue trial. Compared to their peers without stroke, people post-stroke experienced smaller cue-related changes in their BOS-COM distances, as their changes were 1.8 ± 0.6 cm smaller after paretic perturbations with Exact cues (β_P,Ex_ p=0.039) and 2.4 ± 0.6 cm and 1.9 ± 0.6 cm smaller after paretic (β_P,Count_ p=0.002) and non-paretic (β_NP,Count_ p=0.021) perturbations, respectively, with Countdown cues. Overall, people without stroke used cues to bring their COM closer to the edge of the BOS at the recovery step HS, mirroring the MOS results, but people post-stroke maintained similar BOS-COM distances, regardless of the perturbation or cueing condition.

Lastly, we investigated the role of COM speed in cue-related changes to MOS by evaluating changes in fore-aft COM speed at the HS of perturbation (Figure 5A) and recovery steps (Figure 5B). Compared to unperturbed steps, COM speeds during perturbation steps in the No-cue trial were 0.53 ± 0.02 m/s faster for people without stroke (β_NS,No_ p<0.001), 0.32 ± 0.03 m/s faster for people post-stroke on the paretic side (β_P,No_ p<0.001), and 0.33 ± 0.03 m/s faster for people post-stroke on the non-paretic side (β_NP,No_ p<0.001). While there was a significant effect of cueing on the change in COM speed at perturbation step HS (Cue: p=0.041), the effect did not yield significant Bonferroni-corrected comparisons between the No-cue trial and trials with General (β_NS,Gen_ p=0.156), Exact (β_NS,Ex_ p=1.000), or Countdown (β_NS,Count_ p=1.000) cues, and this did not differ between groups (Interaction between Leg and Cue: p=0.074). Although the fore-aft COM speed was faster than unperturbed values at the recovery step HS, there were no effects of group (Leg: p=0.672), cueing (Cue: p=0.158), or the interaction between the two (Interaction between Leg and Cue: p=0.636). Therefore, while treadmill accelerations increased COM speed, as expected, these changes were not mitigated by audiovisual cues.

**Figure 5.**
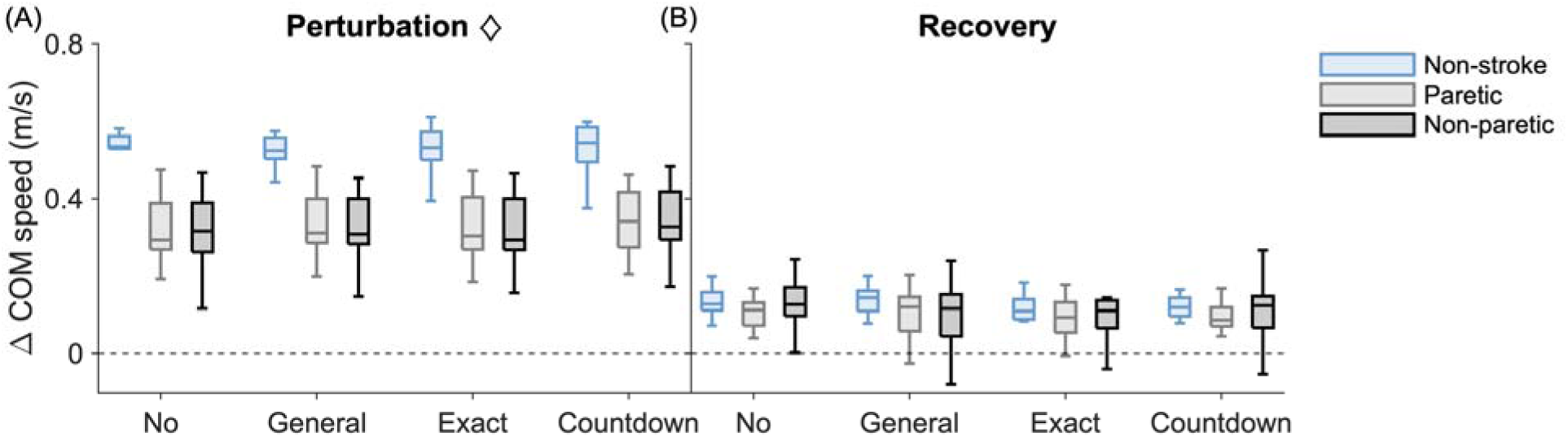
The change in the fore-aft center of mass (COM) speed at heel strike of (A) perturbation and (B) recovery steps with No, General, Exact, and Countdown cues. The changes were calculated relative to unperturbed walking. Boxplot colors represent perturbations to the Non-stroke (blue), Paretic (gray), and Non-paretic (black) limbs. Diamonds in the title indicate significant main effects of Leg. The results of post hoc analyses evaluating the main effect of Cue or its interaction with Leg are represented by vertical gray shading, indicating significant differences from the No-cue trial, and asterisks, indicating significant within-trial differences between the non-stroke and stroke groups. Fourteen total outliers were omitted to maximize the visual size of the boxplots.

### People without stroke performed less positive leg work during cued perturbations

We next evaluated if participants proactively modified positive leg work during the pre-perturbation (Figure 6A) and perturbation (Figure 6B) steps when given cues about impending perturbations. While there was a significant main effect of cueing on changes in positive leg work during the pre-perturbation step (Cue: p=0.015), this effect did not remain after Bonferroni-corrected post hoc comparisons as there were no significant differences in positive leg work between the No-cue trial and the trials with General (β_NS,Gen_ p=1.000), Exact (β_NS,Ex_ p=0.086), or Countdown (β_NS,Count_ p=1.000) cues.

**Figure 6.**
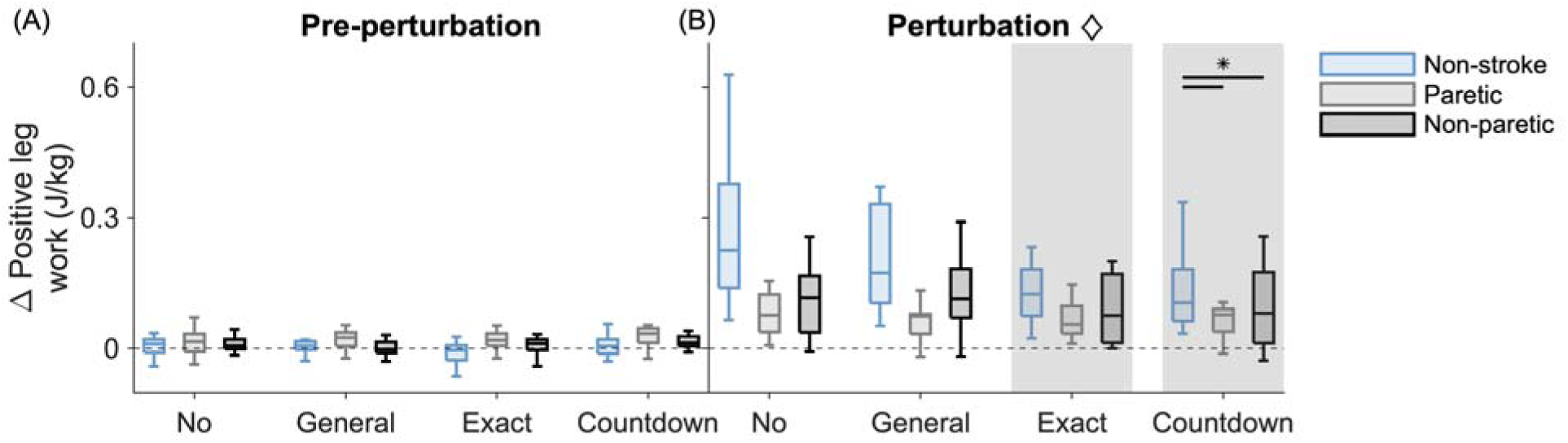
The change in positive leg work during (A) pre-perturbation and (B) perturbation steps with No, General, Exact, and Countdown cues. The changes were calculated relative to unperturbed walking. Boxplot colors represent perturbations to the Non-stroke (blue), Paretic (gray), and Non-paretic (black) limbs. Diamonds in the title indicate significant main effects of Leg. The results of post hoc analyses evaluating the main effect of Cue or its interaction with Leg are represented by vertical gray shading, indicating significant differences from the No-cue trial, and asterisks, indicating significant within-trial differences between the non-stroke and stroke groups. Sixteen total outliers were omitted to maximize the visual size of the boxplots.

During the perturbation step (Figure 6B), the group without stroke performed 0.22 ± 0.02 J/kg more positive leg work than they did on unperturbed steps (β_NS,No_ p<0.001), which was a greater change than the group post-stroke who performed by 0.14 ± 0.03 J/kg during paretic perturbations (β_P,No_ p<0.001) and 0.10 ± 0.03 J/kg during non-paretic perturbations (β_NP,No_ p=0.002). Audiovisual cues also affected changes in positive leg work during the perturbation step, but this change differed between the groups (Cue: p<0.001; Interaction between Leg and Cue: p=0.005). People without stroke performed 0.06 ± 0.01 J/kg and 0.08 ± 0.01 J/kg less positive leg work during perturbations with Exact (β_NS,Ex_ p<0.001) and Countdown (β_NS,Count_ p<0.001) cues, respectively, than during the No-cue trial. Compared to the group without stroke, the group post-stroke made smaller reductions in positive leg work by 0.06 ± 0.02 J/kg and 0.05 ± 0.02 J/kg during paretic (β_P,Count_ p=0.008) and non-paretic (β_NP,Count_ p=0.041) perturbations, respectively, with Countdown cues. Overall, audiovisual cues allowed people, especially those without stroke, to perform less positive leg work during the perturbation steps.

### People without stroke used cues to perform less ankle work and increase BOS-COM distance

Lastly, we evaluated changes in positive ankle, knee, and hip work in the Appendix to better understand the joint-level contributions to leg work changes during the perturbation step. People, especially those without stroke, performed more positive ankle work during perturbed versus unperturbed steps (Appendix Figure 1). When provided any cue, people without stroke were able to perform less positive ankle work during the perturbation step (Appendix Figure 2A). There were minor changes in positive knee work with General and Countdown cues (Appendix Figure 2B) and positive hip work with General cues (Appendix Figure 2C), but the sizes of these changes were smaller than those to ankle work. Cue-related changes to positive ankle (p<0.001) and hip (p=0.045) work were associated with changes in BOS-COM distance at the recovery step HS, with 0.01 J/kg reductions in each corresponding to 0.33 ± 0.04 cm and 0.15 ± 0.07 cm respective increases in BOS-COM (R^2^=0.366).

## Discussion

People post-stroke report frequent falls (12), suggesting their balance control is impaired. Many real-world balance challenges are predictable, allowing for both proactive and reactive control, but little is known about how a stroke influences the roles of each control strategy during predictable gait perturbations. This study investigated how people with and without stroke use audiovisual cues to adjust their balance control strategies during trip-like treadmill perturbations. We found that people without stroke used the Exact and Countdown cues to increase recovery step MOS, and this was associated with a reduction in positive leg and joint work during the perturbations. On the other hand, people post-stroke did not make these proactive adjustments, but instead maintained the same reactive strategies, regardless of cueing, suggesting that control strategies are less tuned to information about impending perturbations in people post-stroke.

### People without stroke use timing cues to reduce the destabilizing effects of treadmill perturbations

Participants without stroke reduced the destabilizing effects of treadmill perturbations when provided audiovisual cues that began one stride (Exact) or three strides (Countdown) before the perturbation. In both conditions, perturbations caused smaller reductions in the subsequent MOS compared to the No-cue trial. In contrast, there were no changes in the MOS with General cues provided three to eight steps before the perturbation, underlining the need for precise information about perturbation timing for participants to proactively update their motor plans. While performance in the Exact and Countdown conditions was similar, future studies will need to explicitly probe if the number of cues and preparation time have independent effects on the use of proactive strategies. Instead, our results suggest that, in some cases, precise timing information in advance of a perturbation can update one’s control strategies to mitigate destabilization.

These findings complement prior work, which demonstrated that people without stroke reduce perturbation-induced increases in COM speed during regular versus random accelerations (29). However, these results differ from other findings that young adults made no significant changes to whole-body angular momentum (WBAM) or MOS during the pre-perturbation and perturbation strides when provided a three-second countdown before treadmill accelerations, decelerations, and waist-pulls (10). It is possible that the treadmill accelerations used in that study were not sufficiently destabilizing or that the young adults felt confident in their reactive strategies, both of which may negate the need to generate proactive adjustments. Notably, the average age of our participants was 58 years old, and we may expect older adults to make more proactive changes if they are less confident in their reactive balance than young adults. Therefore, although the mechanisms by which young adults modulate their proactive control strategies remain unclear, we found that older adults utilize precise audiovisual warnings to improve their stability following treadmill accelerations.

### Modulating mechanical work with cues may be a key proactive control strategy for people without neurologic injury

Next, we investigated the strategies participants used to reduce the destabilizing effects of perturbations when provided with audiovisual cues. We tested for proactive changes during the pre- and perturbation steps and found that people without stroke reduced positive leg and joint work, particularly at the ankle, during the perturbation step, perhaps to slow their forward COM speed or move their COM to a more posterior position during the perturbation. Positive ankle work is generally performed during the push-off phase of stance, so reducing push-off could have helped people improve subsequent stability. Consistent with this interpretation, we found that changes in positive ankle work during the perturbation step were negatively correlated with changes in COM position relative to the edge of the BOS at HS of the recovery step, suggesting people reduced push-off to maintain their COM position. Because changes in forward COM speed were not well explained by changes in joint work, there are likely other strategies, such as foot placement, that play a role in speed control. While changes in positive hip work were also associated with these changes in BOS-COM distance, the effect sizes were larger for the changes in positive ankle work. Parallel changes in positive leg and ankle work are consistent with prior research that found the ankle joint to best reflect changes in leg work during unexpected treadmill accelerations (34). Therefore, performing less positive leg and ankle work during cued perturbations – especially when the cues were more precise – appears to be a key strategy for mitigating the destabilizing effects of predictable perturbations that would otherwise lead to forward losses of balance.

### A stroke may shift dependence onto a general strategy that does not update with cues

In general, we found that people post-stroke did not make proactive balance adjustments when provided with cues about the timing of impending perturbations. Unlike their peers without stroke, people post-stroke did not use the cues to update their proactive strategy and perform less positive leg or ankle work during the perturbation steps on either side. Since the perturbation intensity was comparable across groups with similar changes in fore-aft COM speed, proactive adjustments could still have been beneficial for people post-stroke. As a result, their stability was affected as people post-stroke did not increase their MOS during the recovery step following paretic-side perturbations with Exact or Countdown cues. This group difference is consistent with our previous findings that only people without stroke performed less positive leg work during regular-versus random-interval perturbations on the paretic side (29). It is unclear whether this lack of proactive changes is due to abnormal muscle coordination (27,35) preventing effective changes in muscle activity, spasticity (36) preventing joint extension, sensory dysfunction (20) reducing trust in feedback that could inform proactive updates, or other stroke-related changes to neuromotor control. Future studies should incorporate electromyography, which could characterize anticipatory muscle activity and stroke-related differences in lower-level proactive control strategies. Overall, our findings add to the existing evidence that a stroke impairs proactive balance control strategies during standing (16) and walking (19,18,29).

If proactive control strategies are less flexible and effective following a stroke, a larger reactive response may be necessary to prevent falling. However, a stroke may also impair reactive control during walking, as people post-stroke experience more forward rotation during random treadmill accelerations and require an additional recovery step before returning to baseline than their peers without stroke (14). One reactive strategy may involve adjusting foot placement and COM position to maintain the size of the BOS, whether done passively or actively. Previous research has found strong associations between changes in COM dynamics and subsequent foot placement during walking in both unperturbed (37) and perturbed conditions (38). Here, the group without stroke used the cues to update this proposed control strategy and increase BOS-COM distances on the recovery step, but the group post-stroke did not. Instead, the group post-stroke maintained BOS-COM distances that were similar in magnitude to unperturbed distances, regardless of cueing. This unchanging reactive response, combined with the lack of a proactive change in leg or ankle work, suggests that people post-stroke implement control strategies that are more general and less flexible than those of people without neurologic injury.

Improving balance outcomes is a key rehabilitation goal following a stroke, and the lack of proactive adjustments in our participants post-stroke suggests that future interventions could focus on proactive control. Perturbation training has emerged as a research tool to enable people post-stroke to practice balance control strategies, with some mixed but promising initial improvements in dynamic balance and balance confidence (39–42). However, the emphasis has been on random perturbations that require reactive responses, and it remains unknown whether proactive strategies during gait can be trained and transferred to real-world conditions. If one were to use cued perturbations in mobility training, it is possible that patients may initially rely on preparatory strategies involving a high degree of cognitive control. However, with repeated exposure over multiple sessions, these proactive strategies might become more automatic, increasing the potential functional generalization to real-world walking. Future studies should test this possibility by evaluating if people post-stroke can learn to make effective proactive adjustments such as modulating leg work, ankle work, or step lengths in the presence of predictable perturbations, and determining if these adjustments become more automatic with practice.

### Limitations

There are a few limitations to our current study that could be addressed in future research. We measured changes in MOS, positive leg work, and lower-extremity joint work, but other proactive strategies, such as modulating limb impedance (43–45), may have been implemented to improve stability during cued perturbations. Additionally, although there were no differences in MoCA scores between groups, it remains possible that domain-specific cognitive impairments not captured by the MoCA may have contributed to the lack of proactive changes in the group post-stroke. Future studies should incorporate a comprehensive battery of cognitive assessments to evaluate this possibility. Lastly, because our groups walked at different speeds and the perturbations elicited changes in recovery step MOS with varying magnitudes, it may be challenging to disentangle the effects of stroke versus speed on one’s ability to utilize precise timing cues to improve stability. We scaled the perturbation magnitudes with walking speed to allow people to walk at their comfortable speeds and normalize the perturbations, but it is possible that those walking at slower speeds, often in the group post-stroke, did not experience destabilizations large enough to warrant changes in control. However, our perturbations elicited comparable relative changes in COM speed across groups and conditions, suggesting we perturbed groups similarly. Therefore, while we demonstrated that people post-stroke made limited proactive adjustments during perturbed walking, future studies should continue to investigate the underlying mechanisms of proactive and reactive balance control using other perturbation conditions and tools.

## Conclusions

We aimed to investigate how people with and without stroke modify their proactive control strategies during treadmill perturbations with audiovisual cues. Adults without a history of stroke used audiovisual cues that began one and three strides before treadmill perturbations to mitigate the destabilizing effects, as measured by smaller reductions in the fore-aft margin of stability compared to the margins of stability in the absence of cues. This improvement in stability may have been accomplished by a range of proactive control strategies, including performing less positive leg and joint work, especially at the ankle, during the expected perturbation steps and then better positioning the body’s center of mass relative to the base of support. As hypothesized, people post-stroke made fewer adjustments during predictable perturbations – on both their paretic and non-paretic sides – than their peers without stroke. People post-stroke did not use audiovisual cues to perform less work, but instead, they reactively maintained COM positioning regardless of perturbation predictability. Overall, our findings suggest that people with chronic stroke do not update their control strategy, even when the balance disturbance is explicitly expected, and thus may be at risk for falls that could be prevented with proactive adjustments.

## Appendix

### Supplementary Methods

We characterized changes in positive ankle, knee, and hip joint work during the pre-perturbation and perturbation steps as potential contributors to the proactive changes in positive leg work. First, joint moments were calculated in Visual3D via inverse dynamics, and joint powers were computed in Watts as the dot products of these joint moments and their respective angular velocities. Then, we integrated the positive portions of joint power to evaluate energy generation as the mass-normalized positive work per step in Joules per kilogram. Due to a technical error, joint-level kinetic data were not collected during the Countdown cue trial for one of the participants post-stroke.

We fit separate linear mixed-effect models (Equation 4) for the changes in median positive joint work relative to baseline with fixed effects of Leg, Cue, and the interaction between the two. Specifically evaluating proactive control, we considered changes in positive joint work during the pre-perturbation and perturbation steps relative to unperturbed, baseline walking. If there were significant fixed effects, we reported coefficients and Bonferroni-corrected p-values.

Lastly, we evaluated the relationships between changes in joint work and subsequent changes to two components of MOS at the recovery step HS: the BOS-COM distance or the fore-aft COM speed. Here, we considered the components of MOS independently to determine the role of COM position versus speed in any MOS changes observed with cues. The two linear mixed-effects models included four predictors (the changes in ankle, knee, and hip work from the No-cue trial and Leg) but no intercept.

### Supplementary Results: People without stroke performed less positive joint work during cued perturbations

Along with changes in leg work, we evaluated the changes in ankle, knee, and hip work to probe the joint-level contributions to this balance control strategy. During the pre-perturbation step, there were no changes in positive ankle (Cue: p=0.140; Interaction between Leg and Cue: p=0.739), knee (Cue: p=0.088; Interaction between Leg and Cue: p=0.398), or hip (Cue: p=0.393; Interaction between Leg and Cue: p=0.941). In general, during perturbations in people without stroke, the ankle performs more positive work during push-off (Appendix Figure 1A), the knee performs similar amounts of positive work (Appendix Figure 1B), and the hip performs more positive work (Appendix Figure 1C) compared to unperturbed walking.

**Appendix Figure 1.**
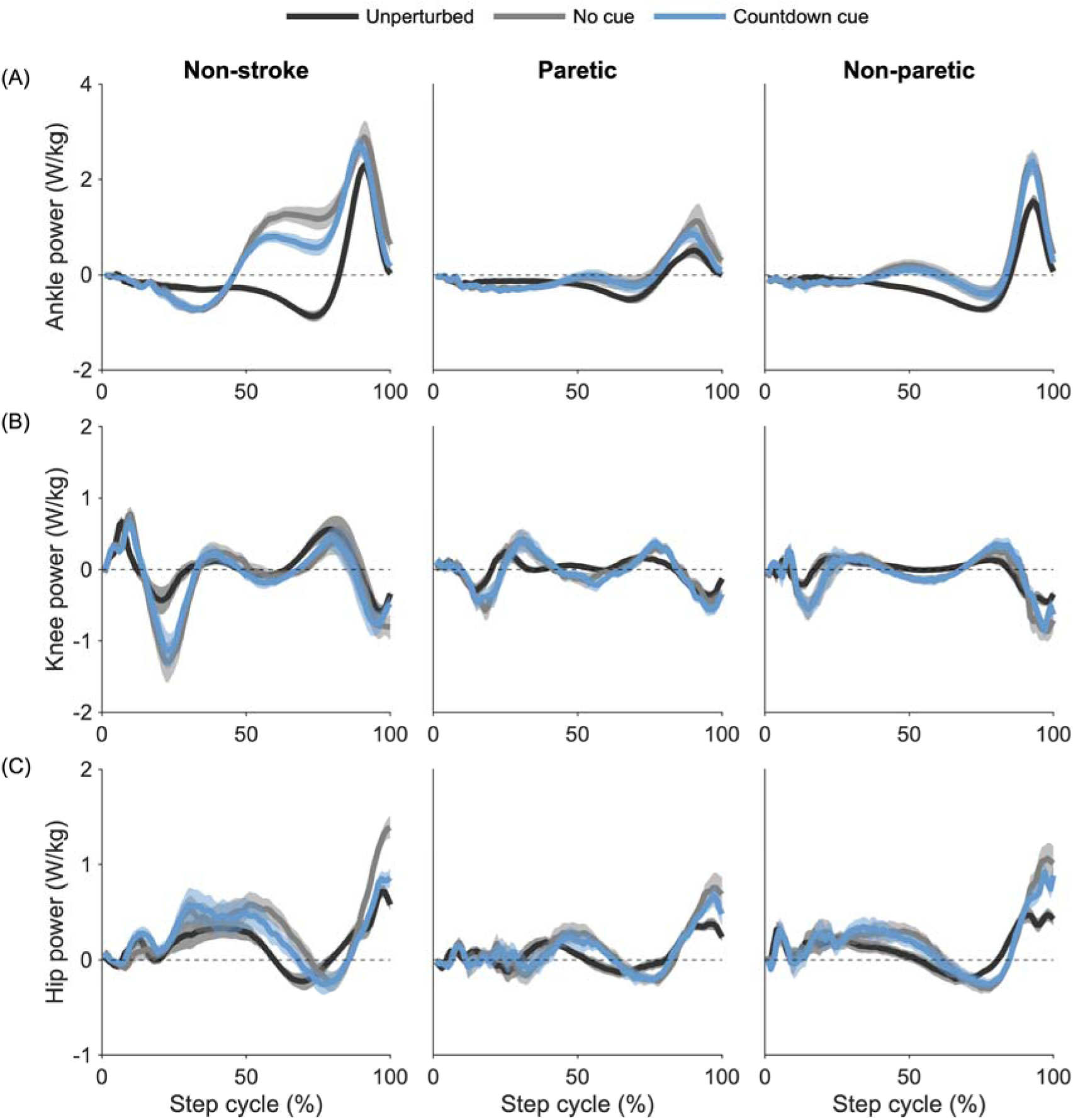
Average (A) ankle, (B) knee, and (C) hip power generated by each leg during unperturbed baseline steps (black), perturbed steps in the No-cue trial (gray), and perturbed steps in the Countdown-cue trial (blue). Trajectories were averaged across strides and then across participants. The ankle performs some negative work during the first half of the stance phase but generates much more positive work during push-off. The knee alternates between performing positive and negative work throughout the step cycle. The hip performs positive work during early to midstance, as well as push-off, and minimal negative work. In general, the paretic limb (middle) performs less positive work relative to the non-paretic limb (right) and to people without stroke (left).

During the perturbation step, people without stroke used cues to perform less positive ankle work. Compared to unperturbed steps, people without stroke performed 0.23 ± 0.01 J/kg more positive ankle work during perturbed steps (Appendix Figure 2A; β_NS,No_ p<0.001). This was a greater increase than the group post-stroke made during paretic (β_P,No_ = -0.17 ± 0.02 J/kg, p<0.001) and non-paretic perturbations (β_NP,No_ = -0.15 ± 0.02 J/kg, p<0.001). Cueing also significantly affected the changes in positive ankle work performed during the perturbation step, though this again differed between groups (Cue p<0.001; Interaction between Leg and Cue: p<0.001). Compared to the No-cue trial, people without stroke performed 0.04 ± 0.01 J/kg, 0.07 ± 0.01 J/kg, and 0.08 ± 0.01 J/kg less positive ankle work during perturbations with General (β_NS,Gen_ p=0.001), Exact (β_NS,Ex_ p<0.001), and Countdown (β_NS,Count_ p<0.001) cues, respectively. However, these cue-related reductions were smaller in the group post-stroke on the paretic side with Exact (β_P,Ex_ = 0.05 ± 0.01 J/kg, p<0.001) and Countdown cues (β_P,Count_ = 0.06 ± 0.01 J/kg, p<0.001) and on the non-paretic side with General (β_NP,Gen_ = 0.04 ± 0.01 J/kg, p=0.046), Exact (β_NP,Ex_ = 0.06 ± 0.01 J/kg, p<0.001), and Countdown (β_NP,Count_ = 0.06 ± 0.01 J/kg, p<0.001) cues.

**Appendix Figure 2.**
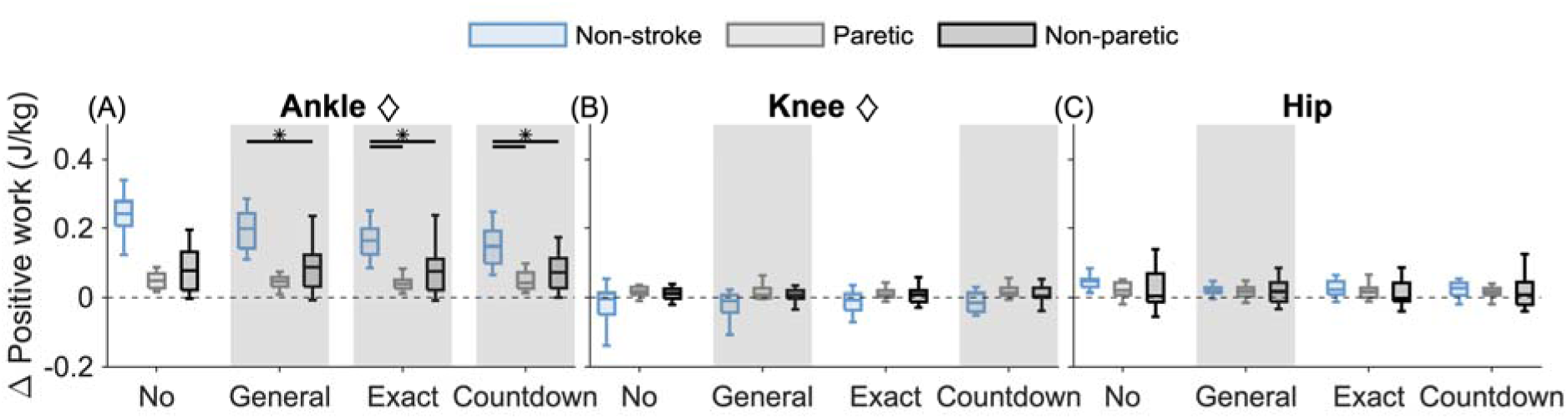
The change in positive (A) ankle, (B) knee, and (C) hip work during perturbation steps with No, General, Exact, and Countdown cues. The changes were calculated relative to unperturbed walking. Boxplot colors represent perturbations to the Non-stroke (blue), Paretic (gray), and Non-paretic (black) limbs. Diamonds in the title represent significant main effects of Leg. The results of post hoc analyses evaluating the main effect of Cue or its interaction with Leg are represented by vertical gray shading, indicating significant differences from the No-cue trial, and asterisks, indicating significant within-trial differences between the non-stroke and stroke groups. Outliers are omitted to maximize the visual size of the boxplots.

We also evaluated if positive knee work changed during perturbation steps (Appendix Figure 2B). While people without stroke did not modify positive knee work during perturbations in the No-cue trial (β_NS,No_ p=0.120), people post-stroke performed 0.04 ± 0.01 J/kg more positive knee work during paretic perturbations than the group without stroke relative to unperturbed values (β_P,No_ p=0.009). There was also a significant effect of cueing on the change in positive knee work during perturbation steps (Cue: p=0.005; Interaction between Leg and Cue: p=0.166), with people performing 0.01 ± 0.00 J/kg less work with General (β_NS,Gen_ p=0.006) and Countdown (β_NS,Count_ p=0.004) cues, but not Exact cues (β_NS,Ex_ p=0.097). While the effect sizes were small, people may have used cues to reduce the amount of positive knee work performed during perturbation steps.

Lastly, we tested for changes in positive hip work during perturbation steps (Appendix Figure 2C). People performed 0.04 ± 0.01 J/kg more positive hip work during perturbed steps in the No-cue trial compared to unperturbed steps (β_NS,No_ p<0.001), and there was no significant difference between groups (Leg: p=0.309). Cueing had a significant effect on positive hip work (Cue: p=0.027; Interaction between Leg and Cue: p=0.264), as people performed 0.02 ± 0.01 J/kg less work with General cues (β_NS,Gen_ p=0.009) but not Exact (β_NS,Ex_ p=0.266) or Countdown (β_NS,Count_ p=0.131) cues. Therefore, people may have used General cues to perform less positive work at the hip during perturbation steps, but the effect size of this change is small relative to the changes at the ankle.

### Supplementary Results: Changes to ankle and hip work are associated with changes to the BOS-COM distance

Finally, we evaluated the relationship between changes in positive joint work during the perturbation steps and each of the components of the recovery step fore-aft MOS: the BOS-COM distance and COM speed at HS. The two models were fit to data collapsed across all cued trials, and fixed effects included the change in positive ankle, knee, and hip work and Leg (Non-stroke, Paretic, Non-paretic). We removed four outliers from the first model and one from the second model before refitting for the final analyses. Changes in positive ankle (p<0.001) and hip (p=0.045) work were associated with changes in BOS-COM distance at the recovery step HS, whereas the change in positive knee work (p=0.684) and Leg (p=0.506) were not (R^2^=0.366). With every 0.01 J/kg reduction in positive ankle work, the BOS-COM distance at the recovery step increased by 0.33 ± 0.04 cm. Additionally, with every 0.01 J/kg reduction in positive hip work, the BOS-COM distance increased by 0.15 ± 0.07 cm. The change in positive knee work (p=0.002), change in positive hip work (p=0.041), and Leg (p=0.018) were associated with the change in COM speed, whereas the change in positive ankle work (p=0.451) was not (R^2^=0.092). However, this model had a poor fit, suggesting that these changes in knee and hip work during cued perturbations were not strongly associated with changes in COM speed.

## Data availability

The processed data and code to create the figures can be found https://osf.io/t8ue4/.

## Acknowledgments

We thank our study participants for their time, effort, and contributions to this work.

## Author contributions

T.C. conceptualized and designed the research, performed experiments, analyzed data, interpreted results of experiments, prepared figures, drafted the manuscript, edited and revised the manuscript, and approved the final version of the manuscript. J.F. conceptualized and designed the research, analyzed data, interpreted results of experiments, edited and revised the manuscript, and approved the final version of the manuscript.

